# Eating in a losing cause: limited benefit of modified macronutrient consumption following infection in the oriental cockroach *Blatta orientalis*

**DOI:** 10.1101/652826

**Authors:** Thorben Sieksmeyer, Shulin He, M. Alejandra Esparza-Mora, Shixiong Jiang, Vesta Petrašiūnaitė, Benno Kuropka, Ronald Banasiak, Mara Jean Julseth, Christoph Weise, Paul R. Johnston, Alejandro Rodríguez-Rojas, Dino P. McMahon

## Abstract

**Background:** Host-pathogen interactions can lead to dramatic changes in host feeding behaviour. One aspect of this includes self-medication, where infected individuals consume substances such as toxins or alter their macronutrient consumption to enhance immune competence. Another widely adopted animal response to infection is illness-induced anorexia, which is thought to assist host immunity directly or by limiting the nutritional resources available to pathogens. Here, we recorded macronutrient preferences of the global pest cockroach, *Blatta orientalis* to investigate how shifts in host macronutrient dietary preference and quantity of carbohydrate (C) and protein (P) interact with immunity following bacterial infection.

**Results:** We find that *B. orientalis* avoids diets enriched for P under normal conditions, and that high P diets reduce cockroach survival in the long term. However, following bacterial challenge, cockroaches significantly reduced their overall nutrient intake, particularly of carbohydrates, and increased the relative ratio of protein (P:C) consumed. Surprisingly, these behavioural shifts had a limited effect on cockroach immunity and survival, with minor changes to immune protein abundance and antimicrobial activity between individuals placed on different diets, regardless of infection status.

**Conclusions:** We show that cockroach feeding behaviour can be modulated by a pathogen, resulting in an illness-induced anorexia-like feeding response and a shift from a C-enriched to a more P:C equal diet. However, our results also indicate that such responses do not provide significant immune protection in *B. orientalis*, suggesting that the host’s dietary shift might also result from random rather than directed behaviour. The lack of an apparent benefit of the shift in feeding behaviour highlights a possible reduced importance for diet in immune regulation in these invasive animals, although further investigations employing pathogens with alternative infection strategies are warranted.

## Background

Microbe symbioses form a fluctuating but universal backdrop to animal life. However, the evolutionary processes that drive animal hosts and their symbionts, including pathogens, operate at different scales and often in opposing directions (Dawkins and Krebs 1979), with the animal immune system acting as a key interface between host and symbiont ecology (Schmid-Hempel 2003). In addition to the core immune system, behavioural mechanisms have attracted increasing attention for their ability to coordinate host responses to infection (Simpson *et al.* 2015; Wong *et al.* 2015). Behaviour is the primary means by which animals interact with the biotic environment, and its importance for a wide range of immune-related functions has recently witnessed a resurgence in research interest.

Hosts can respond behaviourally before infection has even taken place. This can include avoidance of pathogen transmission areas (e.g. defecation sites) and deterrence of disease vectors (Hart 2011; Moore 2013). Other prominent examples include activities falling within the category of ‘social immunity’, which among insects can include pathogen detection alarm behaviours (Rosengaus *et al.* 1999); grooming of conspecific group members (Rosengaus *et al.* 1998; Reber *et al.* 2011); removal (Armitage *et al.* 2016) or even destruction of infected individuals (Yanagawa *et al.* 2011; Davis *et al.* 2018). Such mechanisms are well documented in many social insect lineages, where they contribute significantly to a number of prophylactic mechanisms operating within societies (Schmid-Hempel 1998; Cremer *et al.* 2007). Other prophylactic social behaviours include the collection of secondary antimicrobial compounds to prevent microbial growth in the nest environment (Castella *et al.* 2008; Simone *et al.* 2009), in addition to the direct use – typically via feeding – of antimicrobials in both individual and transgenerational prophylaxis (Lefevre *et al.* 2010; Lefevre *et al.* 2012; Milan *et al.* 2012; de Roode *et al.* 2013; Kacsoh *et al.* 2013).

Once infection has occurred, the first and principal line of defence is the immune system. Here, behavioural defensive adaptations can also play an important role in regulating or augmenting the response to infection. As with prophylaxis, the role of feeding behaviour has increasingly been viewed as a key mechanism by which animals can respond to infection (Abbott 2014). Here, the selection of novel antimicrobial compounds, or the enrichment of specific dietary elements can be employed as therapeutic treatment against pathogens (de Roode *et al.* 2013). Fruit flies use ethanol therapeutically as well as prophylactically to combat parasitoid wasp infection (Milan *et al.* 2012) whereas parasitoid fly-infected *Grammia* caterpillars mix pyrrolizidine alkaloid-producing toxic plants into the normal diet to assist parasitoid clearance, which comes at the expense of body growth (Singer *et al.* 2004; Singer *et al.* 2009; Smilanich *et al.* 2011).

Infection-induced adaptive changes to feeding behaviour can also involve modifications to the quantity and composition of macronutrients in the diet. Anorexia is a well-documented response to infection in both vertebrates (Johnson *et al.* 1993; Konsman *et al.* 2002) and invertebrates (Adamo *et al.* 2007; Ayres and Schneider 2009) and is thought to assist hosts in limiting nutritional resources available to pathogens (Kluger and Rothenburg 1979). Anorexia may also help by activating components of the immune system that are enhanced under conditions of nutritional stress, such as autophagy (van Niekerk *et al.* 2016a; van Niekerk *et al.* 2016b). In recent years, the balance of macronutrients itself has been examined as a way for animals to regulate the response to infection. In particular, the proportion of P has been shown to be an important criterion in animal choice of diet following infection. In *Spodoptera* moths, larvae select a diet enriched in P following infection with a generalist Gram-positive bacterium and a host-specific DNA virus (Lee *et al.* 2006; Povey *et al.* 2009; Povey *et al.* 2013), leading to enhanced antimicrobial activity in both cases. By contrast, diets enriched in C were selected when *Tenebrio* beetles and *Grammia* caterpillars were infected with a rat tapeworm (Ponton *et al.* 2011) and a parasitoid fly (Mason *et al.* 2014), respectively. In the latter study, this behaviour was also associated with an enhanced melanisation response.

The use of macronutrients by hosts to regulate immunity could in principle apply to any animal that is not an obligate food specialist. But less is known about the relationship between macronutrient diet choice and immunity outside of holometabolous insects. Holometabolous insects undergo complete metamorphosis consisting of distinct larval, a pupal and an adult winged phase, which are typically correlated with vastly different ecologies and corresponding physiological, morphological and immunological conditions (McMahon and Hayward 2016). By contrast, hemimetabolous insects undergo progressive molts where each larval instar closely resembles the adult (Sehnal *et al.* 1996). Studies in both locusts and crickets have identified significant correlations between macronutrient intake and immune activity (Rapkin *et al.* 2018; Srygley 2017), but these can result in contrasting effects on host resistance to pathogen infection (Graham *et al.* 2014; Srygley and Jaronski 2018), pointing to a complex relationship between diet, immunity and infection in Orthoptera (Srygley 2016).

Among cockroaches, nutritional studies in *Nauphoeta cinerea* have found that, unlike in other insects (Maklakov *et al.* 2008; Jensen *et al.* 2015a), both sexes prefer a diet enriched in C (Bunning *et al.* 2016). Some studies on the invasive German cockroach, *Blattella germanica*, suggest an apparent robustness to nutritional imbalance and a rapid ability for recovery and dietary adaptation (Raubenheimer and Jones 2006; Shik *et al.* 2014, although see Jensen *et al.* 2015b). This ability may be linked to the fact that cockroaches harbour endosymbiotic bacteria in the fat body that can assist in storing excess nitrogen during over-consumption of P, which can then be redeployed when P is scarce (Sabree *et al.* 2009). Such traits make cockroaches an interesting target for research into the interaction between nutrition and immunity, but this topic has hitherto received relatively little attention.

We tackled this by examining the interaction between macronutrient feeding behaviour and immunity in the omnivorous oriental cockroach, *Blatta orientalis*. We investigated the macronutrient preferences of adult males in response to a range of sublethal immune challenges, before examining the impact of macronutrients on host survival, immune resistance and finally, the expression of the host’s proteome, which captures an additional aspect of the host’s immune response to a pathogen. We tested the hypothesis that *B. orientalis* males modulate macronutrient consumption in response to infection by upregulating the relative intake of P, in turn leading to improved host survival and an enhanced immune response.

## Materials and Methods

### INSECTS AND BACTERIA

A breeding culture of sequential *B. orientalis* cohorts was established at the Federal Institute for Materials Research and Testing (BAM) in June 2015, initially obtained from the collection at the Federal Environment Agency, Berlin, which consists of a mixed population of 4 independent genetic backgrounds maintained for 50 generations. Each generation consists of a minimum of 150 breeding pairs of cockroaches to minimize the effects of inbreeding. Each experimental cohort generation (comprising populations reared independently) was maintained for approximately 190 days in the dark at 26 °C and 50 % humidity, from the day of egg-laying until disposal of older adults. Prior to being placed on experimental (artificial) diets, animals were reared on a mixture of 77.0 % dog biscuit powder, 19.2 % oat flakes and 3.8 % brewer’s yeast and supplied with water *ad libitum* and weekly with apple and carrot slices. All experiments were conducted with adult males (2-3 weeks post eclosion) to minimise changes in physiology associated with oogenesis. Each individual was used only once in each experiment. For the food choice experiment and the survival on enforced diets, individuals from 3 different cohorts were used. The generalist Gram-negative bacterial pathogen *Pseudomonas entomophila* (strain L48; DSM No. 28517) which is able to infect a variety of insect orders (Vallet-Gely *et al.* 2010; Ragheb *et al.* 2017) was obtained from the Leibniz Institute DSMZ-German Collection of Microorganisms and Cell Cultures. Bacteria were stored at −70 °C until use in experiments.

### ARTIFICIAL DIETS

The artificial diets used in this study are based on isocaloric diets, as described elsewhere (Lee *et al.* 2006; Povey *et al.* 2013), which were slightly modified to suit cockroach needs: namely, a drying step was introduced at the end of diet preparation as cockroaches were not able to eat wet food blocks. We employed diets containing 35 % C and 7 % P or *vice versa*, or an equal (E) diet containing 21 % C and 21 % P. The latter diet was selected for some assays because it resembles the composition preferred by cockroaches infected with a high sublethal dose of *P. entomophila* (Fig. 1D) The C portion consisted of sucrose while the P portion consisted of casein, peptone and albumin from eggs in a 3:1:1 ratio. Remaining ingredients are listed in Supplementary Tab. 1. Diet blocks of approximately 0.125 cm³ in size were dried at 50 °C for 2 days before being weighed and given to experimental cockroaches.

**Fig. 1:**
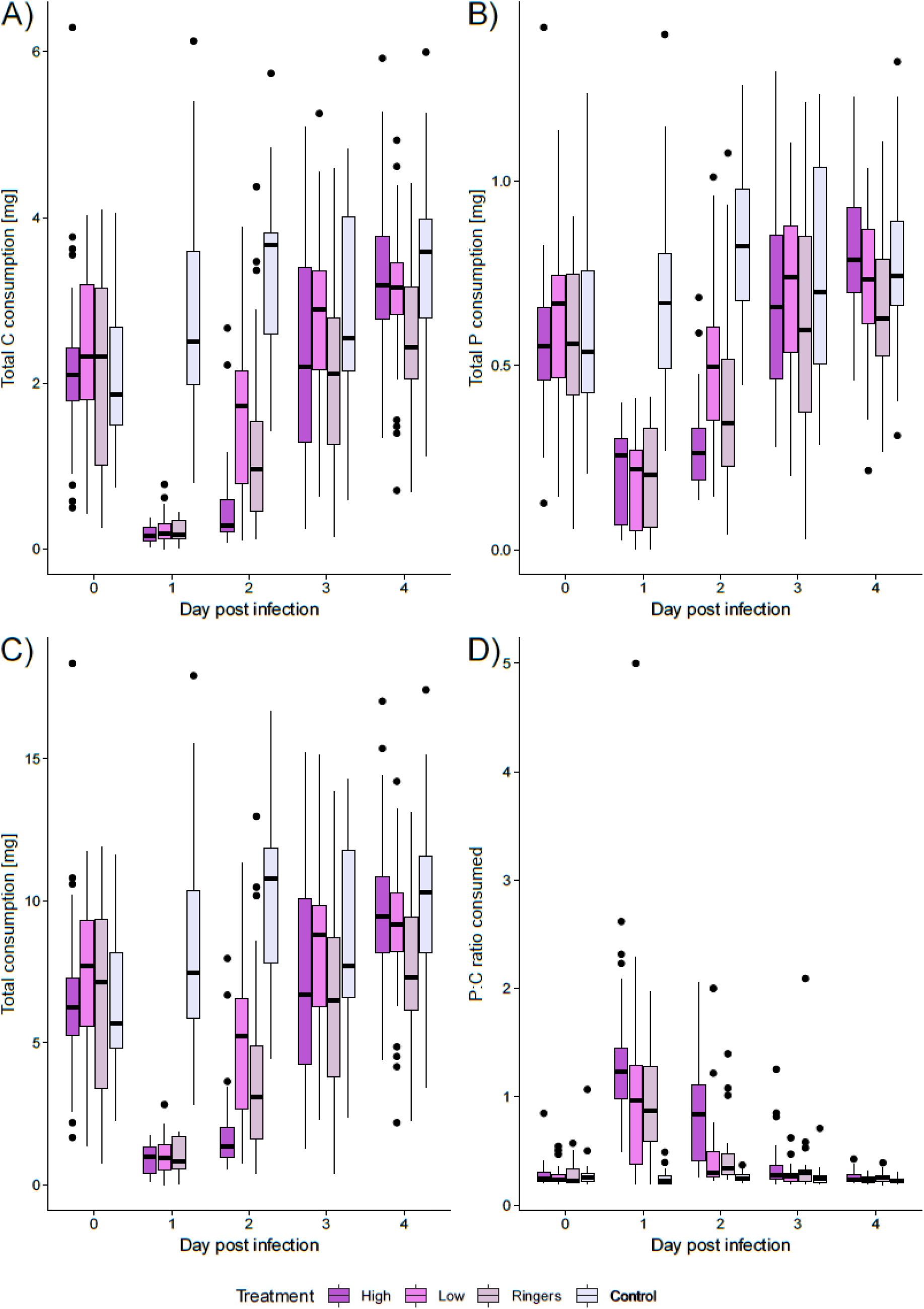
Effect of bacterial infection with *P. entomophila* (high load, low load), Ringer’s solution or no manipulation (control) of *B. orientalis* males on: **A)** C consumption, **B)** P consumption, **C)** total consumption, **D)** P:C ratio consumed. Note different scales used for total P- and C-consumption.

### BACTERIAL INOCULATION

About 200 μl of an overnight culture of *P. entomophila* was mixed in 10 ml fresh liquid medium (according to DSMZ instructions) and incubated at 28 °C and 140 rpm to an OD_600_ of 0.55, representing 1.5 × 10^8^ CFUs per ml. The desired concentrations of bacteria were subsequently obtained by diluting bacteria in insect Ringer’s solution (0.024 g calcium chloride, 0.021 g potassium chloride, 0.01 g sodium hydrogen carbonate, 0.45 g sodium chloride, 200 ml distilled water). Cockroaches were anaesthetised with CO_2_, abdomens swabbed with 70 % ethanol, then injected with 2 μl of bacterial solution directly into the hemocoel using a glass capillary needle inserted between the 3^rd^ and 4^th^ abdominal segment. Sublethal infections (high: 5.8 × 10^5^ CFUs / 2 μl, low: 5.8 × 10^3^ CFUs / 2 μl) and lethal (4.0 × 10^6^ CFUs / 2 μl) doses were determined in pre-experiment injection assays.

### DIET CHOICE FOLLOWING SUBLETHAL INFECTION

From each of 3 cohorts, 40 *B. orientalis* males (120 in total) were given free choice of macronutrients by placing them together with 1 block of known weight of each P-rich and C-rich diet. Individuals were kept for three days to accustom them to artificial diets, and to obtain a baseline P:C ratio preference. Thereafter, food blocks were collected, placed at 50 °C until completely dry, and then their weight loss was determined, equating to the amount eaten by the cockroach. Experimental cockroaches were assigned randomly to one of the following sublethal treatments (40 per treatment): 1) High infection (injected 5.8 × 10^5^ *P. entomophila* CFUs); 2) Low infection (injected 5.8 × 10^3^ *P. entomophila* CFUs); 3) Wounding control (injected Ringer’s solution); 4) Unmanipulated control. Cockroaches were then placed on new food blocks of both diets of known weight. The blocks were replaced daily for four days and their loss of weight was again determined after drying at 50 °C.

### SURVIVAL ON ENFORCED DIET

From each of 3 cohorts, 10 *B. orientalis* males were placed on P-rich diet (35 % P; 7 % C) and another 10 were placed on C-rich diet (7 % P; 35 % C). All individuals were supplied with water *ad libitum*. Survival was checked twice weekly; food blocks and water were changed once a week over the period of 150 days.

### SURVIVAL ON ENFORCED DIET FOLLOWING LETHAL INFECTION

Two hundred and seventy *B. orientalis* males were assigned to one of the following treatments: 1) 150 individuals: Infection (injected 9.0 × 10^5^ *P. entomophila* CFUs); 2) 60 individuals: Wounding control (injected Ringer’s solution); 3) 60 individuals: Unmanipulated control. A third of the individuals from each treatment were randomly assigned to either a P-(35 % P; 7 % C), C-enriched (7 % P; 35 % C) or a U (21 % P; 21 % C) artificial diet and supplied with water *ad libitum*. Survival of each individual was recorded every 2 h for 139 h with overnight intervals of 8 h. The experiment was conducted twice, and the data were combined for subsequent analysis (N=540).

### HEMOLYMPH COLLECTION

Hemolymph for the bacterial growth inhibition assay and proteomic analysis (below) was collected by cutting the first 2 leg pairs of cockroaches which were pre-chilled on ice. They were then placed head-first into a spin-column (Sigma-Aldrich) in a 1.5 ml tube containing propylthiouracil (to inhibit phenol-oxidase activity). They were then centrifuged at 500 *g* for up to 5 min or until at least 10 μl of hemolymph were collected.

### BACTERIA GROWTH INHIBITION ASSAY

In an initial bacterial growth inhibition assay, 180 *B. orientalis* males were equally assigned to the following treatments: 1) bacteria challenge (injected 5.8 × 10^5^ *P. entomophila* CFUs); 2) Wounding control (injected Ringer’s solution); 3) Unmanipulated control. A third of the individuals from each treatment was randomly assigned to either a P-(35 % P; 7 % C), C-enriched (7 % P; 35 % C) or a U (21 % P; 21 % C) artificial diet and supplied with water *ad libitum*. After 24 h the hemolymph of each individual was collected as described in the hemolymph collection section and the hemolymph from 5 individuals per treatment was pooled (resulting in 4 pools per treatment). Pools were stored at −70 °C until needed. A second bacteria growth inhibition assay was conducted on a subset of treatments as an independent validation of the first assay. Methods were identical except 120 *B. orientalis* males were challenged with bacteria (injected 5.8 × 10^5^ *P. entomophila* CFUs) with half of the individuals being randomly assigned to either a P-(35 % P; 7 % C) or C-enriched (7 % P; 35 % C) artificial diet, and hemolymph each from 10 individuals being pooled per treatment (resulting in 6 pools per treatment).

In both assays, bacterial growth inhibition of the cockroach hemolymph was measured using a plate reader assay. First, 10 μl Mueller-Hinton broth were added to each well of a 384-well polypropylene plate. Then 10 μl hemolymph was loaded in the second and the ninth column of the plate. One of these wells contained the hemolymph of one pool of animals (in total 36 wells loaded with hemolymph). Four wells in the first column which did not contain hemolymph served as the negative control. A five-step serial dilution of the hemolymph was performed (with the last 10 μl being discarded) and 10 μl *P. entomophila* in Mueller-Hinton broth with an OD_600_ of 0.005 was added to each well containing hemolymph as well as to another four wells in the ninth column not containing hemolymph, which served as a positive control for unsuppressed bacterial growth. OD_600_ was measured in a plate reader (BioTek) every 10 min for 16 h at room temperature.

### PROTEOMIC ANALYSIS BY MASS SPECTOMETRY

We were unable to detect any significant effect of an equal diet on hemolymph antimicrobial activity, regardless of infection treatment, and so restricted our proteomic analysis to a comparison of the most divergent diets: P-rich versus C-rich following sublethal challenge. One hundred and twenty *B. orientalis* males were immune-challenged by injecting 2 μl Ringer’s solution containing 5.8 × 10^5^ *P. entomophila* CFUs. Half were assigned to the protein-(35 % P; 7 % C) and the other half to the carbohydrate-enriched (7 % P; 35 % C) artificial diet and supplied with water *ad libitum*. Twenty-four hrs later the hemolymph of each individual was collected as described in the hemolymph collection section and stored at −70 °C until needed. This time-point matched the sampling point of the antibacterial assay and was selected to coincide with the peak of infection. The rationale being that for the dietary shift to be relevant for immune activity it must take effect by this point. A detailed description of protein sample preparation and liquid chromatography-mass spectrometry and data processing is described in Supplementary file 1. Initially, we carried out *de novo* transcriptome sequencing to generate a peptide database for *B. orientalis* (Supplementary data sheet 1) Briefly, RNA was extracted from *B. orientalis* by homogenizing individuals in pre-cooled Trizol (Thermo Fisher Scientific) and recovering RNA using chloroform extraction and isopropanol precipitation, and treatment with TurboDNase (Ambion) according to manufacturer’s instructions. mRNA libraries were enriched and prepared using a NEXTflexTM Rapid Directional mRNA-seq Kit protocol (Bioo Scientific) before being sequenced on an Illumina NextSeq500/550 platform at the Berlin Center for Genomics in Biodiversity Research (BeGenDiv). Raw data were processed and annotated as described elsewhere (He *et al.* 2018) (Supplementary file 1). For proteomic analysis, protein identification and label-free quantification was performed using MaxQuant (v1.6.0.1) with Andromeda search engine (Cox and Mann 2008; Cox *et al.* 2011; Tyanova *et al.* 2015). Raw data were matched against an in-house protein database of *B. orientalis* created by *de novo* transcriptome sequencing (see above). Trypsin was selected as enzyme allowing a maximum of two missed cleavages. The minimum peptide length was set to 7 amino acids and the false discovery rate for peptide and protein identification was set to 0.01.

### STATISTICAL ANALYSIS

All statistical analyses were carried out in R v4.0.3 (R Core Team 2020). Testing for normality was performed using the ksnormal function of the wrappedtools package v0.3.11 (Busjahn 2020). P:C ratios, the amounts of P and C eaten as well as total consumption differences between treatments for the first day following infection were analysed using Bonferroni-corrected Wilcoxon rank sum tests.

The food-choice data were analysed using a generalized linear mixed model (GLMM) with an underlying beta family distribution. Analyses were run in the glmmADMB package v0.8.3.3 (Fournier *et al.* 2012; Skaug *et al.* 2014) in conjunction with the R2admb package v0.7.16.2 (Bolker *et al.* 2017). GLMMs examined whether a response variable consisting of proportion of P consumed (amount of P eaten divided by the amount of total diet eaten) or proportion of C (amount of C eaten divided by the amount of total diet eaten) was influenced by treatment (high infection; low infection; wounded; and unmanipulated) and day post infection as well as an interaction between treatment and day. Minimal adequate models were derived by stepwise-model simplification and comparison via ANOVA. Individual and cohort were treated as random effects to account for multiple measurements and origin. Comparisons among treatment levels were carried out with post-hoc Tukey tests using a Bonferroni correction, using package multcomp v1.4-15 (Hothorn *et al.* 2008). Five individuals were removed prior to analysis due to the presence of fungal growth on the artificial diet blocks.

The effect of treatment and diet on survival was analysed using Cox proportional hazard models with the package coxme 2.2-16 (Therneau 2020). Median survival time for each treatment was calculated using the survminer package v0.4.8 (Kassambara *et al.* 2020). Because control data in the survival on enforced diet following infection experiment were right-censored, we uncensored one randomly selected individual from each treatment, following Tragust *et al.* (2013). Owing to the high number of comparisons between treatment levels in this experiment, we conducted post-hoc Tukey tests with Bonferroni or false discovery rate (FDR) corrections and report the results of both methods. Bacterial growth inhibition data were analyzed using R package growthcurver 0.3.1 (Sprouffske 2020) using default parameters. Empirical area under the curve (eAUC) values were analyzed using t-tests in the R package rstatix v0.6.0 (Kassambara 2020). Due to the high number of post hoc comparisons, pairwise t-tests were conducted with Bonferroni as well as FDR corrections. As before, both sets of p-values are reported. In a second analysis bacterial growth inhibition data were combined for a subset of treatments (N=10 replicates per treatment, Pinfected versus Cinfected) from two independent assays, using a two-way ANOVA to examine the eAUC value, with an interaction between treatment and assay.

## Results

### DIET CHOICE FOLLOWING SUBLETHAL INFECTION

Individual cockroaches ate on average 0.56 mg P and 2.24 mg C under unmanipulated conditions, and this remained stable throughout the experiment. Conversely, in all manipulated groups, total food as well as P and C consumption varied significantly over the course of the experiment (Fig. 1A-C). The total amount eaten was reduced in all challenged treatments compared to unmanipulated cockroaches on the first day post-infection (p.i.), but did not differ significantly between manipulated treatments (Wilcoxon rank sum test: high vs. low: W = 429, p > 0.1; high vs. wounded: W = 384.5, p > 0.1; high vs. unmanipulated: W = 0, p < 0.001; low vs. wounded: W = 413.5, p > 0.1; low vs. unmanipulated: W = 0, p < 0.001; wounded vs. unmanipulated: W = 0, p < 0.001). This pattern was replicated in the consumption of P on the first day p.i. (Wilcoxon rank sum test: high vs. low: W = 542.5, p > 0.1; high vs. wounded: W = 452, p > 0.1; high vs. unmanipulated: W = 15, p < 0.001; low vs. wounded: W = 388, p > 0.1; low vs. unmanipulated: W = 12, p < 0.001; wounded vs. unmanipulated: W = 17, p < 0.001) and the consumption of C on the first day p.i. (Wilcoxon rank sum test: high vs. low: W = 382.5, p > 0.1; high vs. wounded: W = 334, p > 0.1; high vs. unmanipulated: W = 0, p < 0.001; low vs. wounded: W = 413.5, p > 0.1; low vs. unmanipulated: W = 0, p < 0.001; wounded vs. unmanipulated: W = 0, p < 0.001).. However, by the 2^nd^ day, consumption across all manipulated groups began to recover, reaching pre-treatment levels by the 4^th^ day p.i..

Before wounding or infection, cockroaches of all treatments preferred a median P:C ratio of approximately 1:4.17 (Fig. 1A). The unmanipulated animals consumed this ratio over the course of the experiment. By contrast, highly infected individuals changed to a P:C ratio of approximately 1.23:1 whereas low infected and wounded cockroaches shifted to an intermediate ratio of 1:1.04 and 1:1.15 P:C on the first day p.i., respectively (Wilcoxon rank sum test: high vs. low: W = 595, p > 0.1; high vs. wounded: W = 571, p > 0.1; high vs. unmanipulated: W = 810, p < 0.001; low vs. wounded: W = 412.5, p > 0.1; low vs. unmanipulated: W = 695.5, p < 0.001; wounded vs. unmanipulated: W = 713.5, p < 0.001). All manipulated groups returned to baseline P:C ratios by day 4 p.i. (Supplementary Tab. 2; Supplementary data sheet 2).

We then carried out GLMMs to explore food consumption differences between treatments over the course of the experiment. Final minimal GLMMs consisted of the fixed terms treatment and day without an interaction since the model with a treatment*day interaction did not significantly improve the model (ANOVA for model comparison, p > 0.1). Cockroaches that were wounded differed significantly from unmanipulated cockroaches in their consumed P proportion following treatment (P proportion chosen: 0.19 vs. 0.08, wounded vs. unmanipulated, respectively: z = −6.132 p < 0.001), as did cockroaches infected with a high (P proportion chosen: 0.23 vs. 0.08, high vs. unmanipulated respectively: z = −13.062, p < 0.001) or low bacterial dose (P proportion chosen: 0.19 vs. 0.08, low vs. unmanipulated respectively: z = −5.332, p < 0.001) (Supplementary Tab. 3; Supplementary Fig. 1). Cockroaches infected with a high bacterial dose also consumed a higher proportion of P compared to individuals exposed to both a low bacterial dose (P proportion chosen: 0.23 vs 0.19, high vs. low respectively: z = −6.258, p < 0.001) or to wounding (P proportion chosen: 0.23 vs. 0.19, high vs. wounded respectively: z = −4.786, p < 0.001). However, individuals that were wounded or were infected with a low bacterial dose did not consume a significantly different proportion of P to each other (P proportion chosen: 0.19 vs. 0.19, low vs. wounded respectively: z = 1.038, p = 1.000). Concerning the proportion of C, the pattern is the same (Supplementary Tab. 3; Supplementary Fig. 2).

### SURVIVAL ON ENFORCED DIET WITHOUT INFECTION

The median (50 %) survival time for *B. orientalis* males placed on a P-rich diet was 82 days, whereas the mortality of males placed on a C-rich diet did not exceed 30 % over the course of the experiment (150 days) (Fig. 2A) (Supplementary data sheet 3). By the end of the experiment males restricted to P-rich diet showed a significantly higher mortality (86.44 %) compared to those on C-rich diet (27.59 %; Cox proportional hazard regression P vs. C: Hazard ratio = 5.73, z = 5.974, *p* < 0.001), although an overt increase in mortality of males restricted to a P-rich diet is only observable after approximately 40 days (Fig. 2A).

**Fig. 2:**
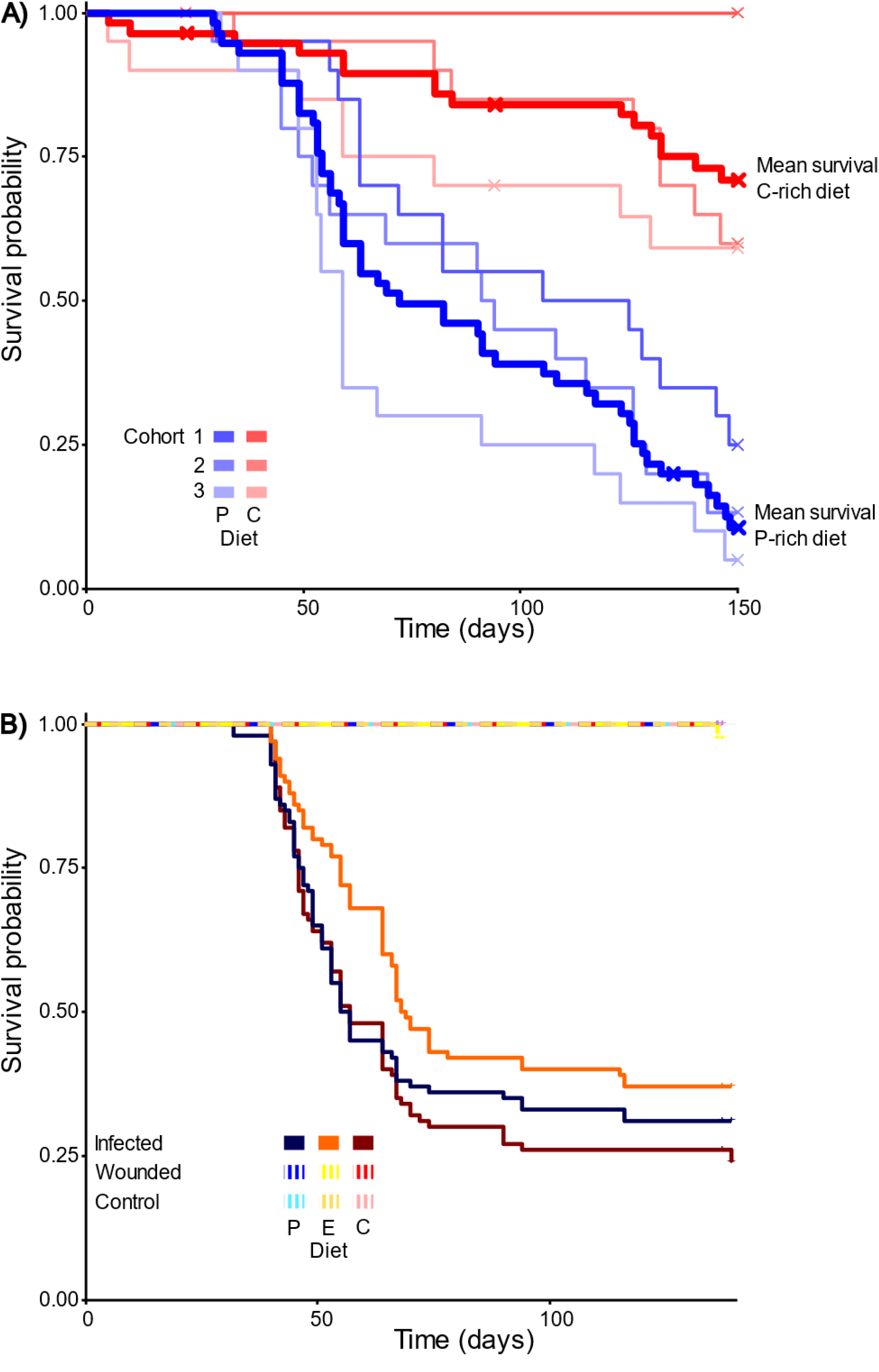
Kaplan-Meier survival curves of: **A)** Unmanipulated *B. orientalis* males restricted to P-rich (35 % protein and 7 % carbohydrate) or C-rich (7 % protein and 35 % carbohydrate) diets. Survival data for three independent cohorts (1-3) for P- and C-rich diets are given in blue and red respectively, with mean population survival across cohorts on each diet indicated by a thick bold line. Note the long period at the beginning of the experiment where no clear survival differences between diets are observable. B) *B. orientalis* males restricted to P-rich (35 % protein and 7 % carbohydrate) (blue line), C-rich (7 % protein and 35 % carbohydrate) (red line, or E (21 % protein and 21 % carbohydrate) (yellow line) diet following injection with an LD50 of *P. entomophila* (infected), Ringer’s solution (wounded) or unmanipulated (control).

### SURVIVAL ON ENFORCED DIET FOLLOWING INFECTION

In our test of the effect of dietary composition on survival following lethal infection, we found that cockroaches on all diets began to die at 40 or 41 hrs after injection (Fig. 2B). This included individuals on the E diet, that is the diet which most closely resembled the ratio consumed by cockroaches following sublethal infection. The median survival time for infected *B. orientalis* males was 56, 57 and 68.5 hrs on P-rich, C-rich and E diets, but the effect of diet on survival following infection was not significant when the Bonferroni correction was implemented (Cox proportional hazard regression: P_infected_ vs. C_infected_: Hazard ratio = 0.89, z = −0.723, *p* = 1.000; E_infected_ vs. C_infected_: Hazard ratio = 0.66, z = 2.504, *p* = 0.443; E_infected_ vs P_infected_: Hazard ratio = 0.73, z = 1.765, *p* = 1.000) (Supplementary data sheet 4, 5; Supplementary Tab. 4). These findings were similar when using FDR, except that survival was significantly higher in infected cockroaches exposed to an E- versus a C-diet (Hazard ratio = 0.66, z = 2.504, p = 0.023) (Supplementary Tab. 5). Only one control individual (wounded, E-diet) died during the course of the experiment.

### BACTERIA GROWTH INHIBITION ASSAY

The inhibitory effect of male *B. orientalis* hemolymph (N = 4 per dilution per treatment) on bacterial growth was not diet-dependent, either within or between treatments (bacteria challenged, wounded or unmanipulated) (Fig. 3) (Supplementary data sheet 6, 7). This was reflected in the non-significant differences of the t-test in the suppression of bacterial growth between all dietary pairwise comparisons, as expressed by eAUC values (P_infected_ vs. C_infected_: t = 0.902, df = 3.212, *p* > 0.1; E_infected_ vs. C_infected_: t = 0.081, df = 5.491, *p* > 0.1; E_infected_ vs. P_infected_: t = 1.085, df = 3.396, *p* > 0.1; P_wounded_ vs. C_wounded_: t = 0.494, df = 3.420, *p* > 0.1; E_wounded_ vs C_wounded_: t = −0.471, df = 4.845, *p* > 0.1; E_wounded_ vs. P_wounded_: t = 1.643, df = 4.177, *p* > 0.1; P_unmanipulated_ vs. C_unmanipulated_: t = 0.332, df = 4.362, *p* > 0.1; E_unmanipulated_ vs. C_unmanipulated_: t = 0.144, df =5.175, *p* > 0.1; E_unmanipulated_ vs. P_unmanipulated_: t = 0.066, df = 3.612, *p* > 0.1.) Some but not all growth curves were significantly different to either the negative or the positive control. No clear pattern was observable between dietary treatments and controls, except that when using a Bonferroni correction, only hemolymph from cockroaches fed on P-diets (bacteria challenged, wounded and unmanipulated), in addition to the negative control, differed significantly from the positive control (Supplementary Tab. 6). Combining a subset of these eAUC values with a supplementary antibacterial assay of P_infected_ vs. C_infected_ (N=10 replicates per treatment) yielded a similarly non-significant result (two-way ANOVA, F = 0.564, df = 1, *p* > 0.1) (Supplementary Tab. 7; Supplementary Fig. 3; Supplementary data sheet 8, 9).

**Fig. 3.**
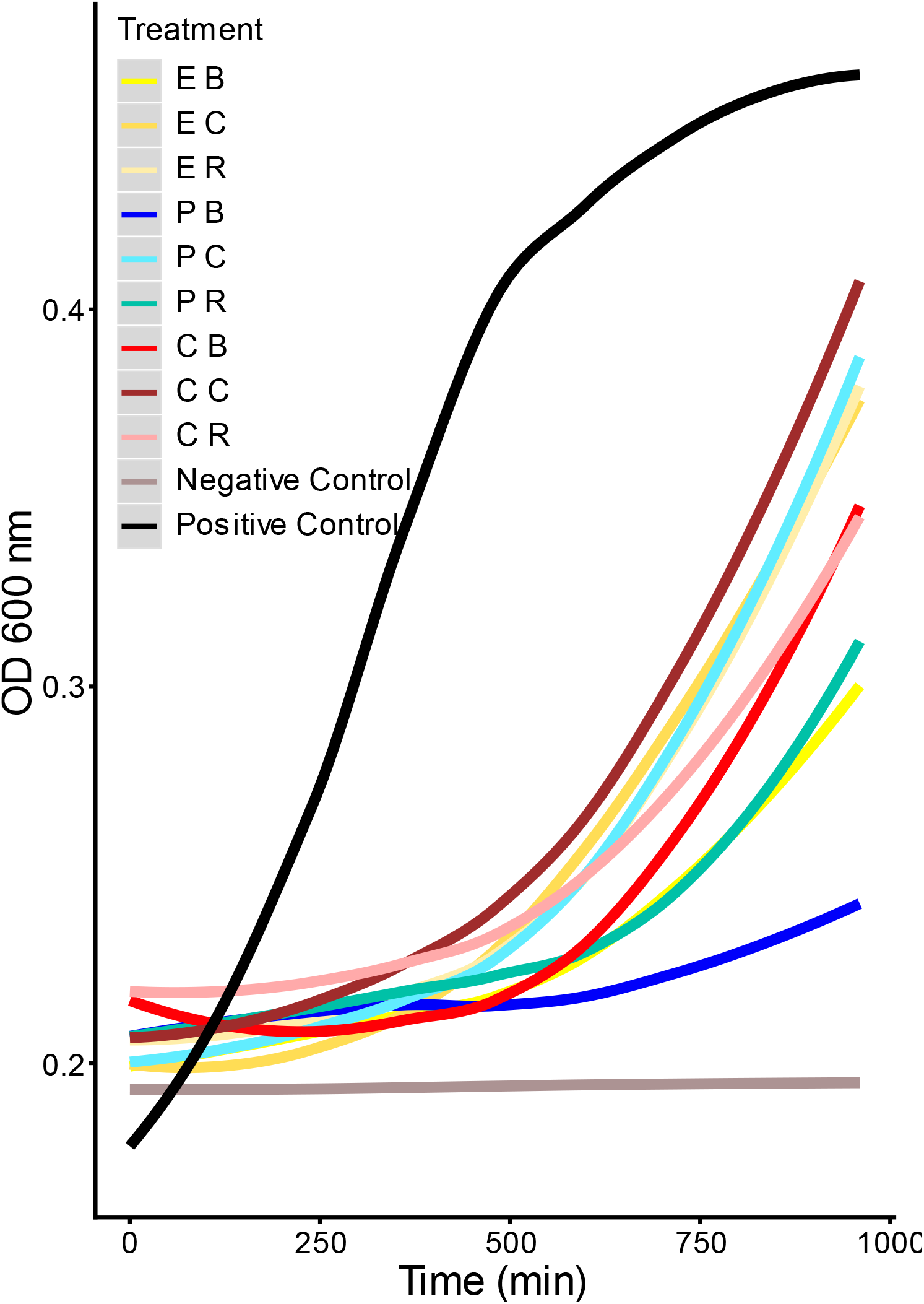
Impact of diet on *B. orientalis* hemolymph growth inhibition of *P. entomophila in vitro* (1:4 dilution). Immune-challenged individuals on P-rich (P B), C-rich (C B) equal (E B) diet. Ringer’s solution injected (wounded) individuals on P-rich (P R), C-rich (C R) or equal (E R) diet. Control (unmanipulated) individuals on P-rich (P C), C-rich (C C) or equal (E C) diet. A bacterial solution without hemolymph served as the positive control and a solution containing only the growth medium (Mueller Hinton) served as the negative control.

### PROTEOMIC ANALYSIS BY MASS SPECTOMETRY

A total number of 3514 peptide hits were identified and assembled into 750 proteins by MaxQuant. After filtering, 387 different proteins were identified and quantified in the hemolymph of infected *B. orientalis* males fed on a P-rich vs. a C-rich diet (N = 6 per treatment) (Supplementary data sheet 10). Overall, apolipophorin was the most abundant protein making up approximately 70 % of the whole hemolymph protein content. Other highly abundant proteins were transferrin, gelsolin, heterochromatin-associated protein MENT and an insulin-like growth factor-binding protein complex. We identified 17 proteins that showed significant changes in abundance following diet treatment (Fig. 5 and Supplementary Tab. 8). Infected individuals on a C-rich diet were significantly enriched for hexokinase type II, which is involved in carbohydrate metabolism (glycolysis) (Yanagawa 1978), in addition to carbonyl reductase I-like, which is involved in NADPH-dependent reduction of active substrates including endogenous and xenobiotic carbonyl compounds (Hoffmann and Maser 2007). Additionally, tropomyosin which is a calcium-dependent regulator of muscle contraction (Pomés *et al.* 2007), and acyl-CoA-binding protein, which carries out lipid-binding transport and suppresses glucose-induced insulin secretion (Færgeman *et al.* 2007; Pasco and Léopold 2012) were more abundant. Furthermore, a L-galactose dehydrogenase-like protein was enriched but its function is not known in insects. Conversely, infected individuals on a P-rich diet were significantly enriched for alpha-amylase, which is involved in carbohydrate metabolism (Terra and Ferreira 1994) and proteasome subunit alpha type-3, which is involved in protein degradation (Rivett 1993). Additionally, hemolymph lipopolysaccharide-binding protein-like (2 isoforms), which binds carbohydrates (foreign particles) (Jomori and Natori 1991) and extracellular superoxide dismutase, which carries out superoxide metabolic processing (Felton and Summers 1995) were detected. Glutamine synthetase is involved in glutamate and glutamine catabolism and biosynthesis (Smartt *et al.* 1998) while adenylate kinase isoenzyme 1 and hexamerin are associated with ATP metabolism (Fujisawa *et al.* 2009) and amino acid and energy storage, respectively (Burmester 1999). There was also an enrichment of ankyrin-1, although its function in insects remains unclear.

## Discussion

Under normal conditions, extensive P consumption shortens the lifespan of many insects including ants, honeybees and flies (Lee *et al.* 2008; Dussutour and Simpson 2009; Fanson *et al.* 2009; Grandison *et al.* 2009; Cook *et al.* 2010; Pirk *et al.* 2010), a finding that is corroborated in our and another study of cockroaches (Hamilton *et al.* 1990). Here, we find that male *B. orientalis* cockroaches showed 45 % higher mortality (Fig. 2A) when restricted to a P- vs. a C-rich diet. One explanation for this consistent observation across study organisms is that elevated levels of P increase TOR signalling. TOR serves as a nutrient sensor linked to macronutrient intake and metabolism, causing a broad anabolic response that is life-shortening over the long term (reviewed in Simpson and Raubenheimer 2009). Other explanations could relate to the toxic effects of breaking down nitrogenous products, and the enhanced production of mitochondrial radical oxygen species, DNA and protein oxidative modifications, membrane fatty acid composition and mitochondrial metabolism (reviewed in Simpson and Raubenheimer 2009). The higher abundance of extracellular superoxide dismutase in cockroach males fed on a P-rich diet (Fig. 4; Supplementary Tab. 8) supports this explanation. Furthermore, the overrepresentation of proteins participating in carbohydrate and protein metabolism in C- vs. P-rich diets, respectively, demonstrate that the diets altered cockroach physiology in the expected direction. For example, the higher abundance of alpha-amylase in the hemolymph of *B. orientalis* males feeding on P-rich diet shows these individuals were metabolizing lower quantities of C. Alpha-amylase is thought to be involved in the breakdown of glycogen, which is the major glucose storage compound in animals. It is employed if not enough C is present in the diet (Mohamed 2004).

**Fig. 4.**
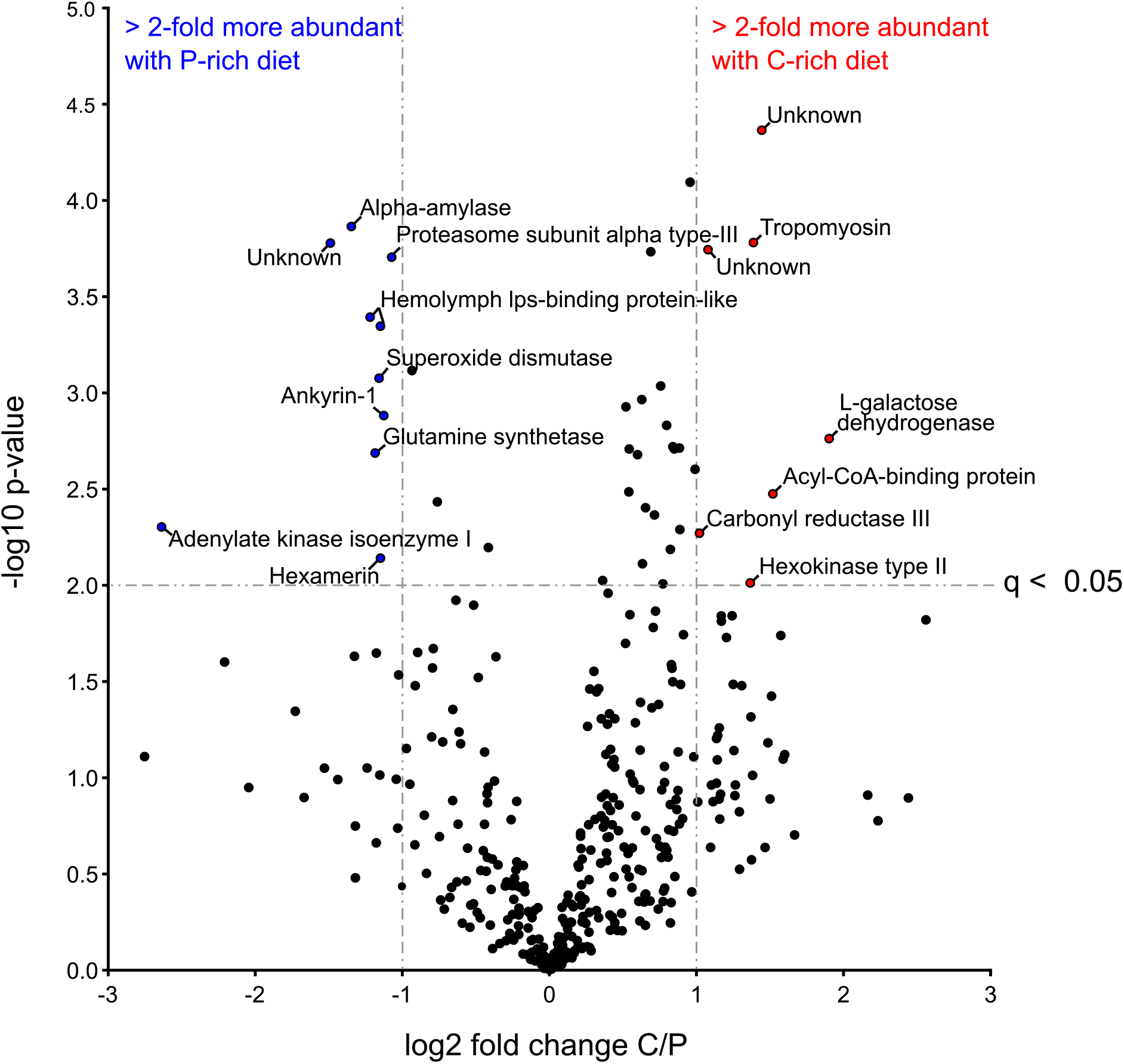
Effect of diet on abundance of male *B. orientalis* hemolymph proteins following bacterial challenge (high dose). Points in blue and red reflect proteins that are significantly (>2) more abundant in P- and C-rich diets respectively.

Unsurprisingly, male cockroaches consumed low amounts of P under normal conditions (1:4.17 P:C). This is in line with the cockroach *N. cinerea*, where males preferred a similarly C-skewed diet of 1:4.8 (P:C) (Bunning *et al.* 2016). Data also indicate that *B. germanica* typically prefers a C-enriched diet, but the degree of C-skew appears to be less pronounced and more variable in this species (Jensen *et al.* 2015b; Jensen and Silverman 2018). In our study, the clear preference for C shifted significantly following infection. As with caterpillars (Povey *et al.* 2013), highly infected male cockroaches increased the ratio of P consumed. Furthermore, cockroaches appeared to adapt their feeding behaviour to the severity of the immune challenge. Lowly infected and wounded (Ringer-injected) individuals consumed an intermediate (approximately uniform) P:C ratio and their food consumption returned to normal sooner after challenge compared to highly infected individuals, which shifted to the most P-enriched diet and displayed the longest delay in returning to normal dietary consumption. It is interesting to note that the observed feeding responses were transient across all challenge treatments, with most individuals returning to a normal dietary intake 72-96 hours after injection. Transience is likely correlated with the period of acute bacterial infection, although wounding itself also elicited a similar response to lowly infected individuals, suggesting a generalized precautionary host response to challenge. Together, our data indicate that *B. orientalis* males are able to quantitatively regulate their behavioural response to infection and rapidly return to a normal feeding regime. Additionally, our findings suggest host-driven adaptation as opposed to pathogen manipulation because wounded individuals also reduced their C intake. Wounding elicits a localized immune response in insects (Haine *et al.* 2007), suggesting a form of prophylactic behaviour since it is likely that microbes can enter the hemolymph via damaged cuticle (Siva-Jothy *et al.* 2005).

In contrast to *Spodoptera exempta* caterpillars and other organisms which can modulate their immune response with diet, changes in cockroach dietary consumption following infection did not greatly influence any of the immune parameters we measured. In caterpillars, a shift from a C- to a P-biased diet following *Bacillus subtilis* (Gram-positive) or baculovirus infection led to an increase of antibacterial and phenol-oxidase activity and hemocyte density and resulted in higher survival (Povey *et al.* 2009; Povey *et al.* 2013). By contrast, a switch to a protein enriched diet did not have a major influence on male *B. orientalis* hemolymph antimicrobial activity or survival, nor have a substantial impact on the synthesis of induced immune-related proteins. We note, however, that two hemolymph lipopolysaccharide-binding protein isoforms, which may play a role in pathogen recognition by binding foreign particles (Jomori and Natori 1991) were more abundant in the hemolymph of P-rich fed infected cockroaches. Furthermore, we observed some evidence for reduced bacterial growth in hemolymph extracted from P-fed cockroaches (compared to the positive control), suggesting that dietary protein may confer some inhibitory effect on bacterial proliferation. However, this effect was not observed with the E-diet (the dietary blend consumed by males following infection), nor was this pattern consistently observed across post-hoc methods. Furthermore, no significant effect of diet was observed between any of the challenge treatments. Interestingly, we found that survival following infection was significantly higher in cockroaches fed on an E- versus a C-diet but again, this effect was not consistent across correction methods, and was not corroborated by other dietary comparisons, as might be expected (e.g. E- versus P-diet, or P-versus C-diet).

Overall, our findings suggest that a shift to a protein enriched diet could have a minor influence on *B. orientalis* immunity, but that in general, the behavioural changes adopted by this cockroach following direct injection with *P. entomophila* are unable to substantially alter infection outcome. However, the longer-term consequences of P on *B. orientalis* immunity, including over the course of development, remain to be investigated. Entomopathogenic pathogens that act more slowly on the host should also be examined in this context. Studies in Orthoptera indicate that an enforced P-rich diet can enhance immune activity both over ontological time and in the short term. However, the benefits of P for host survival after infection are conflicting and appear to depend significantly on host and pathogen identity (Graham *et al.* 2014; Srygley and Jaronski 2018). Graham *et al.* (2014) found that locusts feeding on a C-enriched diet were more resistant to fungal infection, even though enforced consumption of P enhanced several immune parameters, including antibacterial activity. But this outcome may be due to the metabolic requirements of the pathogen in question (Srygley and Jaronski 2018). With respect to host physiology, a recent study suggests that a diet enriched in P may protect against infection simply via the modulation of hemolymph osmolarity (Wilson *et al.* 2020). A potential hypothesis to explore here would be whether natural variation in hemolymph osmolarity might explain the relative (in)effectiveness of P dietary manipulation in different insects following bacterial infection. It is interesting to note that hemolymph osmolarity may be higher in cockroaches compared with other insects, including lepidopterans (Natochin and Parnova 1987), although this requires much further testing. Similarly, it would be important to explore the feeding shifts of a greater diversity of hemimetabolous groups, inclusive of cockroaches, under challenge from a range of pathogenic microbes to understand whether the patterns we observed in *B. orientalis* are associated with specific adaptations such as extreme omnivory and/or endosymbiosis.

Taken together, our results suggest that *B. orientalis* males may not be able to effectively self-medicate against *P. entomophila* using macronutrients, but that they do engage in a typical anorexia response, as has been shown in macronutrient self-medication in caterpillars (Adamo *et al.* 2007; Povey *et al.* 2009; Povey *et al.* 2013). Illness-induced anorexia offsets physiological trade-offs between launching immune responses and food digestion. A previous study demonstrated that crickets reduce their food intake, especially for lipids, following infection with the bacterium *Serratia marcescens* (Adamo *et al.* 2010). High hemolymph lipid levels are associated with decreased concentrations of monomeric apolipophorin III, a lipid transporter, and higher susceptibility to *S. marcescens* infection (Adamo *et al.* 2008). In other insects, anorexia can have a direct impact on immunity. For example, in *Drosophila*, starvation can modify AMP production and lead to reduced melanisation (Ayres and Schneider 2009).

The apparent lack of a link between macronutrient dietary selection and male cockroach immunity is unexpected. One possible explanation is that future food availability and quality may be less predictable in omnivorous pest organisms like cockroaches (Raubenheimer and Jones 2006). A recent genomic study reports major expansions of cockroach gene families linked to chemoreception, detoxification and innate immunity (Li *et al.* 2018), indicating that adaptations in these pathways permit cockroaches to thrive in unpredictable, antigen-rich environments. Indeed, while cockroach survival was reduced on an enforced P-rich diet, a negative effect could only be observed well over 40 days after exposure, suggesting that although protein is generally avoided by *B. orientalis* adult males, it’s consumption can be tolerated for long periods of time. Cockroaches are known to tolerate high levels of P consumption (Cochran 1985) and in such extreme omnivores, there could be an advantage to reducing regulatory interactions between host diet and immunity. This ability could be mediated by the presence of the endosymbiont *Blattabacterium*, which may also be able to help the host store and catalyze excess nitrogen (Patiño-Navarrete *et al.* 2014). An additional point to consider is that in contrast to several previous studies, we performed experiments on adult individuals and not larvae, which have different resource allocation strategies and consumption rates in general (Boggs 2009). Particularly in holometabolous insects, most of the resources in larvae are allocated to growth, maintenance and storage whereas in adults, they are allocated to reproduction and maintenance. Consequently, there has been a greater emphasis on a trade-off between growth and immunity in the larval stage of herbivorous insects (reviewed in Singer *et al.* 2014). The need for fast growth could compete with the requirement to provide protection from parasite and pathogen-induced mortality. Given that P and amino acids are a crucial limiting factor in herbivorous diets, we hypothesize that a trade-off between two essential life-history parameters that depend strongly on P – growth and survival – could be more pronounced in herbivorous juvenile insects (Schoonhoven *et al.* 2005; Simpson and Raubenheimer 2012).

## Conclusion

We find that *B. orientalis* males modulate their macronutrient feeding behaviour following infection by dramatically reducing food intake and simultaneously reducing carbohydrate over protein intake. We also show that a P-rich diet eventually leads to significantly reduced host lifespan, and that male cockroaches prefer a C-rich diet under normal conditions. To our surprise, the observed behavioural response to immune challenge did not meaningfully influence the antimicrobial activity or proteomic profile of host immunity. Our findings therefore support the concept of a generalized host-directed response to microbial challenge in cockroaches based on anorexia and the limitation of C intake. In this scenario, the observed change to a more equal ratio of P:C may instead reflect a shift towards a severely reduced baseline level of random feeding rather than a directed shift towards a higher ratio of consumed P, although this hypothesis requires additional testing. Such a response may be beneficial to the host, but perhaps primarily as a means of avoiding contaminated food and reducing pathogen access to resources, rather than facilitating crosstalk with the immune system. From an evolutionary perspective, this could be the result of adaptations to detoxification, endosymbiont-mediated metabolism and innate immunity combining to enhance cockroach survival in antigen-rich and nutritionally diverse environments. Overall, our study highlights the importance of understanding variation in natural diet, development, and ecology when exploring the link between nutrition and animal immunity.

## Declarations

### Ethics approval and consent to participate

No ethical guidance or approval was required for working with *Blatta orientalis*.

### Consent for publication

Not applicable

### Availability of data and materials

The short read data used to generate the *Blatta orientalis* transcriptome are available on the SRA (ID: SRX8891863, part of Bioproject PRJNA635910). All other data generated or analyzed during this study are included in this published article and its supplementary information files.

### Competing interests

The authors declare that they have no competing interests.

### Funding

S.H. was supported by the Chinese Scholarship Council and D.P.M. was supported by a seed-funding grant provided by the Freie Universität Berlin and grant MC 436/6-1 from the Deutsche Forschungsgemeinschaft (DFG). We also acknowledge assistance of the Core Facility BioSupraMol supported by the DFG.

### Authors’ contributions

DPM conceived and coordinated the study; TS designed and conducted the experiments; TS and DPM wrote the manuscript; SH and PRJ conducted the proteomic database preparation by de novo transcriptome sequencing; MAE-M, SJ, VP and MJJ assisted the survival and antimicrobial assays; ARR guided the sample preparation for the proteomic analysis; BK and CW conducted LC-MS/MS and corresponding data analysis. All authors contributed to drafting and revising the manuscript.

## Acknowledgements

We thank J. Rolff for providing useful advice and technical support.

## Notes

### Competing Interest Statement

The authors have declared no competing interest.

